# Beeporter: a high-throughput tool for non-invasive analysis of honey bee virus infection

**DOI:** 10.1101/2021.01.13.426606

**Authors:** Jay D. Evans, Olubukola Banmeke, Evan C. Palmer-Young, Yanping Chen, Eugene V. Ryabov

**Author notes:** E-mail addresses,.

## Abstract

Honey bees face numerous pests and pathogens but arguably none are as devastating as Deformed wing virus (DWV). Development of antiviral therapeutics and virus-resistant honey bee lines to control DWV in honey bees is slowed by the lack of a cost-effective high-throughput screening of DWV infection. Currently, analysis of virus infection and screening for antiviral treatments in bees and their colonies is tedious, requiring a well-equipped molecular biology laboratory and the use of hazardous chemicals. Here we utilize a cDNA clone of DWV tagged with green fluorescent protein (GFP) to develop the Beeporter assay, a method for detection and quantification of DWV infection in live honey bees. The assay involves infection of honey bee pupae by injecting a standardized DWV-GFP inoculum, followed by incubation for up to 44 hours. GFP fluorescence is recorded at intervals via commonly available long-wave UV light sources and a smartphone camera or a standard ultraviolet transilluminator gel imaging system. Nonlethal DWV monitoring allows high-throughput screening of antiviral candidates and a direct breeding tool for identifying honey bee parents with increased antivirus resistance. For even more rapid drug screening, we also describe a method for screening bees using 96-well trays and a spectrophotometer.

## INTRODUCTION

Insects and other arthropods are prime vectors of disease agents, including a suite of viruses important for human, plant, and animal health. In addition, many insects play beneficial roles in nature and for humankind. Pollination by insects adds at least $20 billion annually to the U.S. agricultural economy [1] and hundreds of billions of dollars worldwide [2]. Honey bees are the preeminent agricultural pollinators’, thanks to their numbers and the mobility of their colonies. Despite their critical roles in agriculture through pollination and hive products’, honey bee populations are under threat. Half of all honey bee colonies are lost and replaced annually in the U.S. [3]. Along with the financial costs of colony replacement, these losses impact the abilities of beekeepers to fulfil contracts to pollinate almonds and other high-value crops.

Diseases are a major cause of colony losses, especially diseases that are driven by parasitic mites. While mites themselves impact bees, it is the viruses they carry, and Deformed wing virus (DWV) in particular, that drive colony mortality [4, 5]. Nevertheless, no effective treatments are commercially available for honey bee viruses. Identification of new viral controls requires the screening of hundreds of candidate drugs, in line with similar efforts for human medicine and large-animal disease work. This high-throughput screening is best carried out with living bees since this allows the simultaneous identification of off-target effects of chemicals on bees themselves.

A number of cloned viruses expressing fluorescent reporters, mainly mammalian and plant, have been developed [6-8]. These viruses have been used to assess virus replication in a range of cells, tissues, and organisms, greatly advancing our understanding of virus biology and virus-host interactions. Here we describe a protocol for *in vivo* monitoring of viral growth in honey bees. The strength of this protocol is the pairing of infectious viral clones with a gene encoding a fluorescent reporter [9] linked with a virulent US strain of this virus [10]. Bees infected by these clones provide a reliable visual signal that can be screened readily and quantitatively with a standard digital camera. By comparing fluorescent signals with quantitative-RT-PCR estimates of viral loads in the same bees, we see excellent correlations, indicating that GFP fluorescence by itself is a good surrogate for molecular quantification of viral load. The monitoring of DWV infection by observing (recording) GFP fluorescence in live pupae allows investigations of viral replication dynamics at the level of insects. In order to increase sensitivity and specificity, especially in adult bees where darkened cuticles can obscure fluorescence, we also describe an assay reliant on fluorescence measurements using a plate-reader spectrophotometer. After optimizing each strategy, we present the methods used for validation and refinement of conditions.

Honey bee researchers can benefit from this system via virus assays that would normally involve flying bees or bees in cages, saving labor costs. The fluorescent reporter allows for immediate non-invasive screens of viral loads, without expensive and time-consuming RNA extraction and quantitative PCR steps. Companies and researchers seeking new bee medicines will be able to study in-house or novel candidate drugs against these viruses, speeding their searches and reducing the need for specialists in molecular biology [11]. Regulators devoted to testing the impacts of pesticides and other stressors on bee health could assess those impacts through virus loads via this method since viruses are a strong indicator of honey bee stress [12].

Finally, honey bee queen breeding is a million-dollar industry. Bee breeders seeking viral resistance can benefit from a reliable and quick assay for viral resistance in different bee lineages. Breeders could incorporate this system in their selective breeding program, giving them an integrated estimate of virus resistance in their breeder lines, without molecular-genetic resources or skills. In fact, since one of the two protocols described below is non-lethal, individual queens and reproductive males in specific breeding programs might be screened during development as a direct assay for their own breeding value prior to mating. This latter trait has not previously been possible since current screening tools for honey bee viruses involve sacrificing bees for RNA extraction. Further, honey bee males are haploid and effectively all cells in their bodies, including sperm, are genetically identical. Therefore, screens for the abilities of male honey bees to resist diseases provide an especially potent method for shaping breeding populations, since male phenotypes are traceable to single alleles. Haploid males have been exploited to identify resistance traits against parasitic mites [13] but their power in identifying resistance to pathogens has yet to be tapped. Instrumental insemination is used routinely to produce high-value breeder queens and sperm from male bees with confirmed virus resistance can be collected and used directly in bee crosses.

In summary, while genetic techniques are available for quantifying virus loads, these techniques are time consuming, relatively expensive, lethal to subjects, and dependent on expensive fixed laboratory equipment. This protocol allows high-throughput *in vivo* assays of drugs and hosts’ immune defenses. It is highly flexible in terms of host life stages or tissues and, given investment in the development of infectious clones, is applicable in all insect-virus systems.

## MATERIALS AND METHODS

### Reagents and Biological Materials

- Strong honey bee source colonies maintained with low levels (below 1%) of the parasitic *Varroa destructor* mites that vector Deformed wing virus.
- Full-length DWV cDNA clone derived from virulent Maryland isolate pDWV-304 (GenBank accession number MG831200) with sequence encoding enhanced green fluorescent protein, eGFP, or another fluorescent reporter) inserted in frame at the leader protein-viral protein 2 (LP-VP2) border of the DWV cDNA [9].
- 0.22 μm nylon membrane syringe filter (Thermofisher)
- Disposable (100 mm) Petri dishes
- Whatman filter paper folded to separate injected pupae
- Phosphate buffered saline (PBS)
- Disposable 31G insulin syringes, with 6mm needle, capacity 300 7L (BD Becton Dickison)

#### Optional materials, for assay validation, not required for protocol

- Black, clear-bottom 96-well plates (Corning)
- For quantification of DWV loads in the honey bee by RT-qPCR: TRIzol reagent, RNeasy RNA extraction kit (optional), Superscript III Reverse transcriptase, random hexanucleotides, SYBR-Green mix (and DWV- and DWV-GFP-specific qPCR primers as in [9].

### Equipment

- Incubator with manual (dish) system for humidity control
- Micro-syringe injection pump and a microprocessor-based controller, UMP3/Micro4 (WPI - World Precision Instruments).
- UV transillumination table with low-wavelength (395 nm) light source and/or handheld UV ‘blacklight at 365 nm.
- Digital smartphone camera. Can also use gel-documentation system with attached camera (UVP, Bioimaging Systems)

#### Optional equipment, for assay validation, not required for protocol

- Fluorescence-equipped plate reader capable of excitation at 480 nm and measurement of emission at 520 nm, SpectraMax Paradigm™ Multi-mode detection platform (Molecular Devices, LLC, San Jose, CA), for assay validation, not required for protocol.
- Bio-Rad CFX-400 optical thermal cycler, for validation of new assays, not needed for protocol.

### Production of inocula of the virus expressing green fluorescent protein (GFP) reporter

1. Construct or acquire infectious cDNA DWV-GFP plasmid clone (Ryabov et al 2020).
2. Produce the full-length DWV-GFP *in vitro* RNA transcript from linearized plasmid DNA template by HiScribe T7 RNA polymerase (New England Biolabs) according to manufacturer’s guidance, remove plasmid DNA template by treating with Turbo DNAse (Ambion), subsequently subject to phenol-chloroform, chloroform extraction, precipitate RNA by mixing with 2.5 volumes of ethanol and 0.1 volume of 3 M sodium acetate pH 5.2, and precipitating at 12.000 g for 15 min using microcentrifuge, wash visible pellet with 80% ethanol, air-dry, dissolve in RNAse-free water, quantify concentration using Nanodrop and store in -80°C.
3. Inject white-eye honey bee pupae with 5 μg (13.2 × 10^12^ copies) of *in vitro* RNA transcript suspended in 8 uL of PBS.
4. Incubate pupae for 72 hours (hr) at +33°C and 85% relative humidity.
5. Homogenize infected pupal tissue in 1 mL of PBS
6. Subject to three rounds of freeze-thawing and filter through a 0.22 µm nylon filter. Typically, 1 mL of the filtered extract prepared from a single transcript-injected pupae will contain 10^9^ to 10^10^ viral genome equivalents (GE), sufficient for 1000 to 10,000 pupal injections. Note. Filtered DWV-GFP inocula could be produced in equipped molecular laboratory, stored at -80°C for at least 7 months and then shipped in dry ice to end users.

### Injection and incubation of honey bees

7. Collect frames of sealed honey bee worker brood from colonies with low levels of the mite *Varroa destructor*, vector for DWV.
8. Harvest early stage (white-eyed or pale pink-eyed) pupae from comb brood cells, remove carefully to avoid damage, make sure that selected pupae are not *Varroa-*infested.
9. Incubate collected pupae on Whatman paper (Supplemental Figure 1A). in Petri dishes for 3 hr +33°C and 85% relative humidity to detect and dispose pupae damaged during extraction showing development of melanization.
10. Prepare working solutions containing viral inoculum in PBS (generally 12 x 10^5^ or 12 x 10^6^ GE of cloned DWV-GFP per µl in for the current trials.
11. For drug testing, dissolve candidates into inocula (generally in a final solution from 1 to 10 ppm) immediately prior to injection.
12. Inject each pupa intra-abdominally, dorsal-laterally (Supplementary Fig. 1B) with 8 µl of viral inocula (with or without drugs) or PBS control using disposable insulin syringe with 31G needle, the same syringe load (typically 240 µL) could be used to inject up to 30 bees.
13. Incubate pupae at +33°C and 85% relative humidity (Fig. 1B).

**Figure 1.**
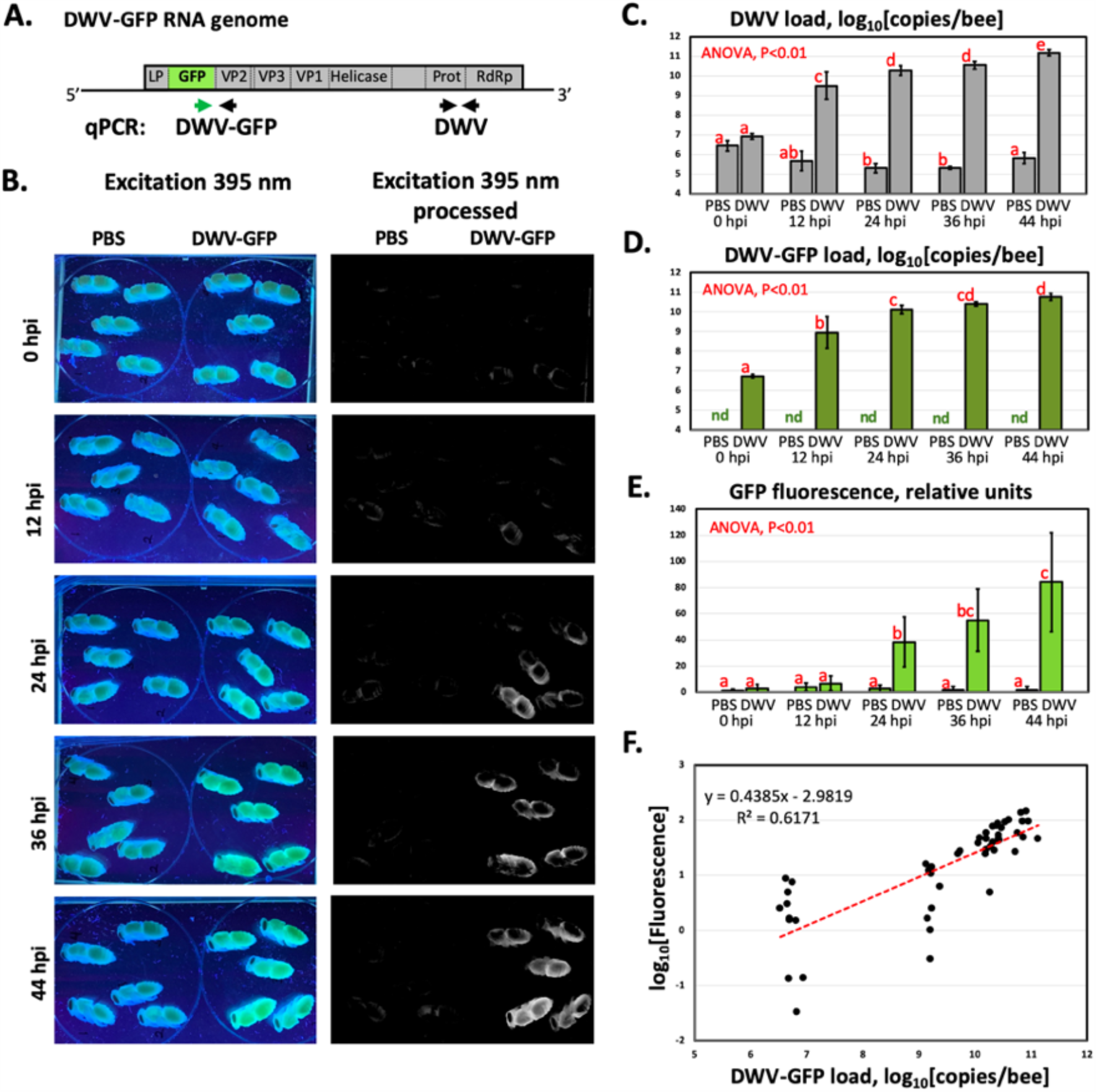
A) Schematic diagram of the DWV-GFP construct (10.8 kb) and the positions of the RT-qPCR primers: “DWV” - generic DWV primers, “DWV-GFP” – tag-specific primers, and B) Successive images of bees from time of injection then 12, 24, 36, and 44 hours later, 395 nm excitation with and without filtering for autofluorescence. For each time panel, left five bees are PBS-injected control, right five – injected with 10^6^ GE of DWV-GFP. C) Quantified viral loads using generic DWV primers. D) Quantified DWV-GFP virus using tag-specific primers. E) Quantified GFP fluorescence over time. F) Log_10_ transformed DWV-GFP loads (RTqPCR quantification) versus GFP fluorescence (log_10_ transformed), with best-fit regression.

### Fluorescence measurements for living bees

14. Acquire digital image of pupae at 0, 12, 24, 36, and 44 hours post injection (hpi) using a standard ultraviolet light table (365 nm wavelength). Each image should include PBS-injected control pupae and/or an adjacent fluorescent standard to normalize images taken on different dates with a handheld smartphone digital camera. Each image should include PBS-injected control pupae and/or an adjacent fluorescent standard to normalize images taken on different dates.
15. For UV gel imaging transilluminator (365 nm excitation) and camera use “SYBR-green” option adjust exposure time and contrast to minimize non-specific background fluorescence in the control PBS-injected pupae (Fig. 3B).

Note: 365 nm and 395 nm UV light sources are used in inexpensive and widely commercially available as “UV counterfeit currency detectors”.

16. Analyze images using ImageJ (https://imagej.nih.gov) [14] for average and maximum green fluorescence of each of five pupae in each treatment group at each time point.

### Fluorescence measurements using a 96-well plate reader

17. Inject honey bee individuals with the tagged virus, 10^6^ genome copies and incubate for 44 hours.
18. Homogenize the pupa with 300 µL of PBS supplemented with a protease inhibitor cocktail (cOmplete, Roche) in 1.5 mL Eppendorf tube on ice. Subject to one cycle of freeze-thaw, spin in a table-top micro centrifuge, 5,000 per minutes for 5 minutes. Collect 100 □L of supernatant for fluorimetry, freeze the rest at –80°C for RNA extraction using TRIzol for subsequent quantification of DWV-GFP loads by RT-qPCR [15].
19. Autofluorescence of pupal tissue by long-wavelength UV light presents the main challenge for visualization of fluorescence of GFP, which was expressed from the viral genome in honey bee tissues. To optimize sensitivity and specificity of the GFP fluorescence detection in honeybee pupal extracts, contrast control and infected bees along a continuum of emission wavelengths (490 nm to 600 nm) after establishing ideal excitation wavelengths (blanketed by 360 nm to 510 nm). This analysis showed that for overt levels of DWV infection, 10^10^ to 10^12^ GE per insect, optimal excitation and emission wavelengths are 480 nm and 520 nm respectively (Fig. 2B). Importantly, this analysis showed that it was possible to use non-optimal excitation UV wavelengths, 365 nm and 395 nm, which are readily available in UV transilluminators (Fig. 2B, pointed with arrows).
20. Measure emittance at optimal conditions (520 nm emission after excitation at 480 nm for the stages we tested).
21. Analyze differential emittance for GFP-carrying bees from various genetic lineages or treatments versus control samples.

**Figure 2.**
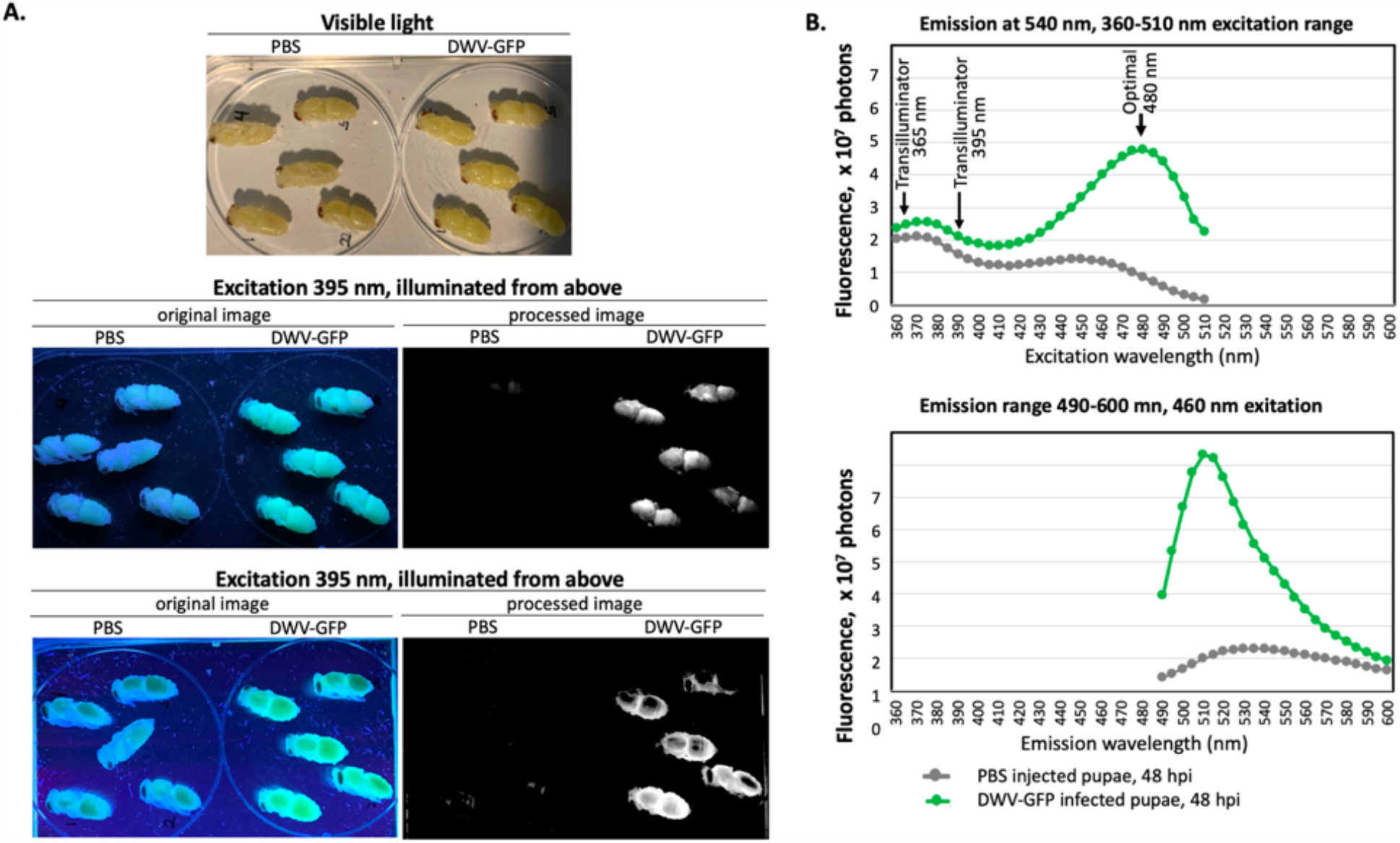
A) Visible, original ultraviolet (365 nm excitation), and filtered ultraviolet images of bees used to test fluorimetry assay. The DWV-GFP infected bees (44 hpi) contained 7.2 x 10^10^ to 9.2 x 10^10^ genome equivalents of DWV-GFP. B) Range of excitation and emissions values (average) used to pinpoint the most accurate and sensitive conditions for the assay. Arrows indicate excitation wavelengths used in UV transilluminators (365 nm and 395 nm), and the optimal excitation wavelength (480 nm).

**Figure 3.**
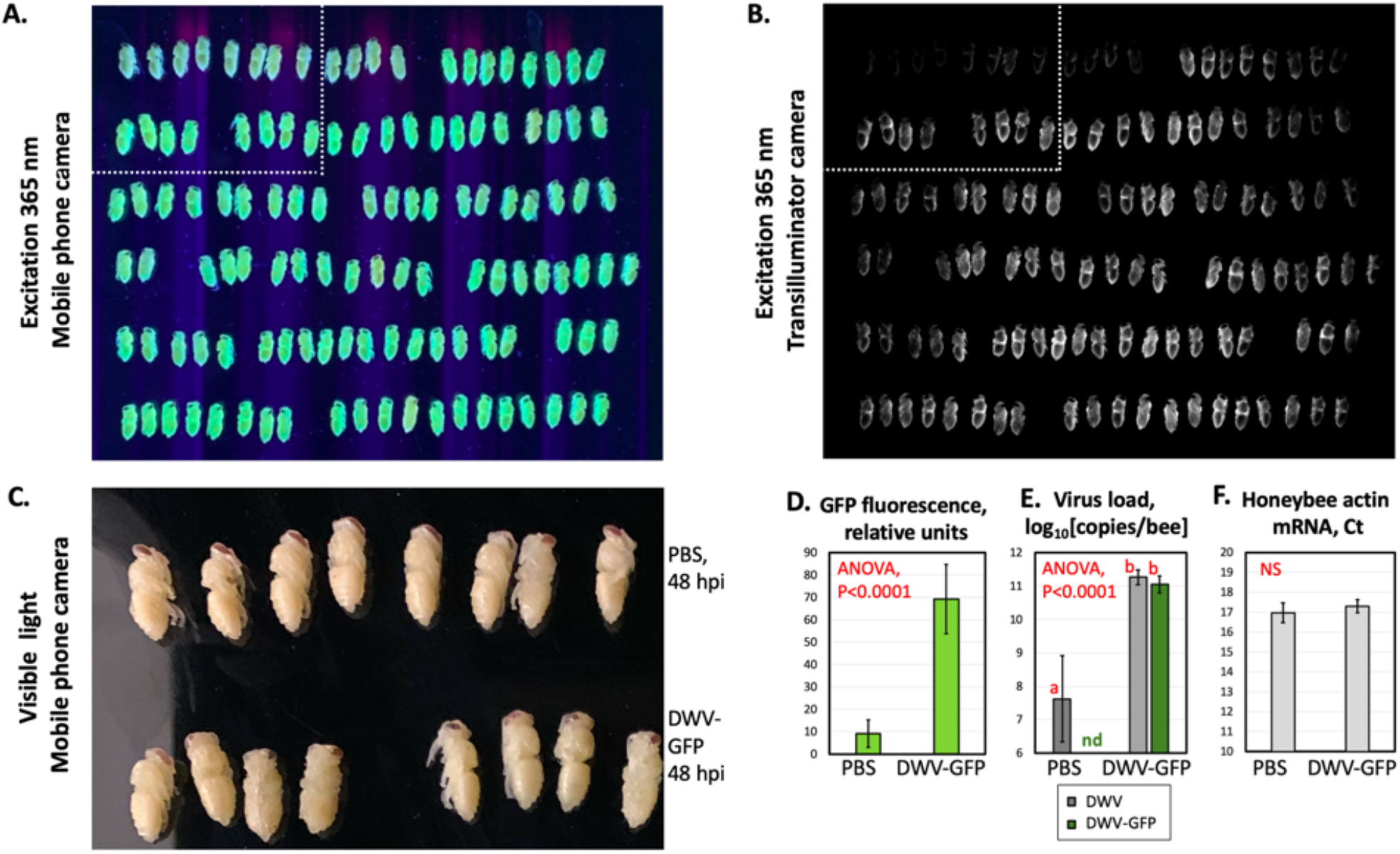
Typical simultaneous visualization of DWV infection in 120 bees, 48 hours post injection (hpi) using standard 365 nm UV transilluminator. Top left 12 bees are injected with PBS control, the rest were injected with 10^6^ copies of DWV-GFP. A. Unprocessed mobile phone camera image. B. Processed gel recording camera, SYBR-green recording option. C. Visible light photograph of a subset of bees (marked in B and C), mobile phone camera image. D-F. Analysis of the PBS and DWV-GFP-injected bees in the marked areas of A and B. D. GFP fluorescence quantified using image B, E, quantification of DWV by RT-qPCR, GE per bee, DWV-specific primers; F, estimation of honey bee actin mRNA loads, Ct values.

### Validation and refinement using quantitative RT-PCR

22. After measuring fluorescence, extract total RNA from samples using the TriZOL method.
23. Measure viral loads using qPCR and validated primers [9].
24. Use PCR thresholds (CT values) for a dilution series of pDWV-GFP plasmid from 10^2^ to 10^8^ copies to establish an efficiency curve for the quantitative PCR runs (y = -0.2666x + 10.829, R^2^ = 0.9889, in our trials).
25. Directly quantify DWV genome copies in the samples and compare log-transformed qPCR values and estimated fluorescence using regression and ANOVA.

### Troubleshooting and critical points

When choosing pupae for injection (Steps 3, 7), use honey bee colonies with low Varroa mite infestation, which are likely to have low levels of Varroa mite vectored wild type DWV. For precise single-virus studies is advisable to monitor colonies for the presence of other honey bee viruses common for the region. Each newly prepared batch of filtered DWV-GFP inoculum (Step 6) should be analysed by RT-qPCR to determine copy numbers of GFP and DWV RNA. Choose the filtered extracts with 1:1 ratios of GFP to DWV copy numbers which has minimal proportion of the clone-derived virus with deletion of GFP. When using DWV-GFP inoculum produced by an external provider, make sure that it was delivered frozen, keep at -80°C or use immediately, keep the inocula on ice. Avoid freeze thawing more than three times.

### Time required

One skilled technician can inject approximately 120 bees/hour and prep the same number of bees for photographic capture in 30 minutes, followed by two hours of statistical analyses using ImageJ [14]. Injecting pupae can also be carried out via manual syringe or Micro-syringe injection pump. To quantify viral loads by fluorimetry post-experiment requires approximately four hours for 120 bees, from pulverizing bees to refinement and capture using the fluorimeter.

## RESULTS

Average fluorescence of bees injected with GFP-tagged DWV or the PBS control placed on the UV transilluminator, excitation wavelengths 395 nm (Fig. 1B, 2A) or 365 nm (Figure 3A,B), was recorded using images taken with an unaided smartphone camera (Fig. 1B, 2A, 3A) or by UV gel imager camera, SYBR green set up (Fig. 3B). Filtration post-hoc using ImageJ greatly reduced autofluorescence. Fluorescence was significantly higher in infected bees versus those injected with the PBS control starting from 24 hpi, and continued to increase at 36 and 44 hpi (Fig. 1B,E, Fig. 3D), as was the estimated viral titer using qPCR (Fig. 1C,D, Fig. 3E). Note that although background levels of wild-type DWV were present even in the PBS-injected bees at low of 10^6^ to 10^7^ GE (Fig. 1C, PBS), only the injected DWV-GFP replicated to high levels (Fig. 1 C, D), which was in agreement with the results of previously reported clone-derived DWV injection experiments [10]. The use of DWV-GFP-specific pair of qPCR primers (Fig. 1A) allowed to detect exclusively the injected clone-derived virus (Fig. 1D).

Quantitative RT-PCR estimates of DWV-GFP (Fig. 1C) and DWV-GFP (Fig. 1D) were strongly correlated with fluorescence estimates (Figure 1F; r^2^ = 0.617). As expected, a linear dependence was observed between the log-transformed DWV-GFP copy numbers quantified by qRT-PCR and log-transformed GFP fluorescence. With a fluorometric plate reader, it was possible to refine further the optimal excitation and emissions wavelengths for this assay (Fig. 2). Ultimately conditions were established with virtually no masking or autofluorescence by host bee tissues, even for a range of bee life stages.

## DISCUSSION

The described methods have general usefulness for tracking insect-virus dynamics. From mosquitoes and other vectors of medical and veterinary importance to aphids, moths, and beetles attacking plants, being able to track viral loads in real time offers a method for resolving how and when viruses proliferate in their hosts. This can predict vectoring ability and better define the stages of viral infection that impact host behavior. In honey bees, these protocols provide a critical tool for high-throughput screening of potential antiviral or host-enhancing drugs [11], and a further refinement of efforts to use functional genetics to better understand bee/virus interactions [9, 16]. This nonlethal method also has considerable promise for bee breeding, since labelled individuals, once affirmed as being resistant to virus infection, can be used directly in breeding efforts. Finally, the impacts of insect viruses on host development and behavior are poorly understood. A system for tracking viral spread and abundance in live insects should prove useful for determining more precisely when infection becomes pathology.

## ACKNOWLEDGEMENTS

This work was supported by the USDA - Agricultural Research Service, ARS project number 0500-00093-001-00-D, USDA National Institute of Food and Agriculture grant 2017-06481 and USDA-APHIS Interagency Agreement 18-8130-0787-IA.

## SUPPLEMENTAL DATA

**Supplementary Figure 1.**
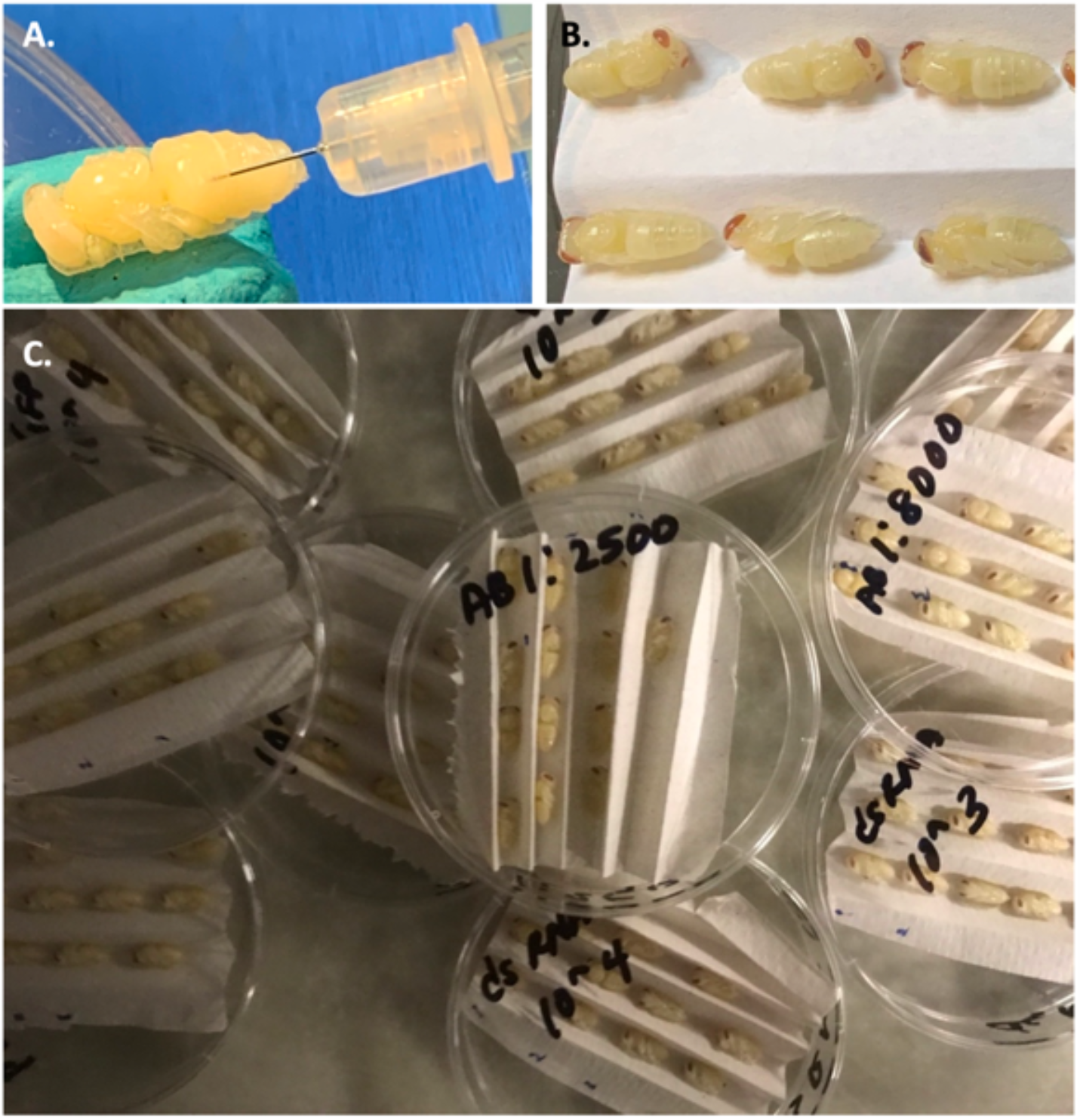
Visual display of bee injection experiment. A. Injection with a 31-gauge needle syringe. B, C. Injected bees. Simultaneous visualization of DWV infection, 48 hours post injection (hpi) in 120 bees using standard 365 nm UV transilluminator, top left 12 bees are injected with PBS control. A. Bees illuminated with 365 nm UV light, unprocessed mobile phone camera image. B. Bees illuminated with 365 nm UV light, processed gel recording camera, SYBR-green recording option (Model??). C. Visible light photograph of a subset of bees (marked in B and C), mobile phone camera image. D-F. Analysis of the PBS and DWV-GFP-injected bees in the marked areas of A and B: D - GFP fluorescence quantified using image B; E – quantification of DWV by RT-qPCR, GE per bee, DWV-specific primers; F – estimation of honeybee actin mRNA loads, Ct values.

